# Elevated and sustained reactive oxygen species levels facilitate mesoderm formation during early *Xenopus* development

**DOI:** 10.1101/223453

**Authors:** Yue Han, Yaoyao Chen, Nick R. Love, Shoko Ishibashi, Enrique Amaya

**Author notes:** Other authors’ email addresses: Yue Han, Yaoyao Chen, Nick R. Love, Shoko Ishibashi.

## Abstract

Fertilisation triggers embryonic development culminating with the activation of a number of highly co-ordinated and evolutionarily conserved signalling pathways, which induce and pattern the mesoderm of the developing embryo. Previous studies in invertebrates have shown that hydrogen peroxide (H_2_O_2_), a reactive oxygen species (ROS), can act as a signalling molecule for axis specification during early development. Using a HyPer transgenic *Xenopus laevis* line that expresses a H_2_O_2_-sensitive fluorescent protein sensor maternally, we recently found that fertilisation triggers a rapid increase in ROS production. Here we show that this increase in ROS levels is sustained throughout early embryogenesis, lasting until the tailbud stages. In addition we show that lowering ROS levels from the blastula stage through the gastrula stages via antioxidant treatments disrupts mesoderm formation. Furthermore, we show that attenuating ROS levels during the blastula / gastrula stages affects some, but not all, growth factor signalling pathways involved in mesoderm induction and patterning, including the PI3K/Akt, TGF-β/Nodal, and Wnt/β-catenin signalling pathways. These data suggest that sustained elevated ROS levels during the blastula and gastrula stages are essential for early vertebrate embryonic development, at least partly, through their roles in promoting growth factor signalling.

## Introduction

More than a century ago, Otto Warburg showed that fertilisation is accompanied by an increase in oxygen consumption in sea urchin embryos (Shearer, 1922; Warburg, 1908). Many decades later, it was shown that this dramatic increase in oxygen consumption was attributed to a burst of reactive oxygen species (ROS) production, including H_2_O_2_ (Foerder et al., 1978). This “respiratory burst” of H_2_O_2_ in sea urchins is generated by an NADPH oxidase, Udx1, and the primary role for this oxidative burst is to cross-link proteins that harden the fertilisation envelope, thus preventing polyspermy (Heinecke and Shapiro, 1989; Wong et al., 2004).

Cellular ROS production has several sources, but the primary ones are NADPH oxidases (NOXes) and the mitochondrial electron transport chain, which generate superoxide (O_2_^δ∂-^), which is rapidly converted to H_2_O_2_, either spontaneously or via the enzymatic activity of superoxide dismutases (Brand, 2016) (Bedard and Krause, 2007). Recent studies have shown that H_2_O_2_ contributes to cellular signalling by modulating a variety of redox sensitive proteins involved in signal transduction. In sea urchin embryos, mitochondrial H_2_O_2_ has been shown to regulate the oral–aboral axis specification by activating the nodal signalling pathway (Coffman et al., 2009; 2014). Recently, Gauron et. al showed that H_2_O_2_ participates in axonal projections via its ability to modulate Shh signalling in zebrafish larvae (Gauron et al., 2016). However, a role of ROS production during early vertebrate development remains largely unexplored.

Embryonic development involves a variety of cellular processes, including cell growth, proliferation, differentiation, migration and death. The precise regulation of these events is essential for morphogenesis and development. A number of highly interconnected and evolutionarily conserved growth factor signalling pathways play critical roles during early embryogenesis, including the induction and patterning of the germ layers (Houston, 2017; Tseng et al., 2017). Previous studies have shown that exogenously or endogenously produced ROS can inhibit or activate various signalling pathways, such as MAPK signalling (Kamata et al., 2005; Robinson et al., 1999; Santabárbara-Ruiz et al., 2015) (Ray et al., 2012), PI3K signalling (Cho et al., 2004), Wnt/β-catenin signalling (Funato et al., 2006), nodal signalling (Coffman et al., 2014; 2009) and Hedgehog signalling (Gauron et al., 2016). We previously showed that sustained ROS production is necessary for appendage regeneration in *Xenopus* tadpoles, in part via their ability to promote FGF and Wnt/β-catenin signalling (Love et al., 2013). Given that both FGF and Wnt/β-catenin signalling also play essential roles during mesoderm formation and patterning during embryogenesis, we wondered whether ROS might also play critical roles during mesoderm formation in early embryos.

Using a previously established transgenic line, HyPer (Love et al., 2013; 2011), which allows the assessment of ROS levels *in vivo,* we recently showed that fertilisation triggers a burst of ROS production in *Xenopus,* which plays a critical role in the regulation of the early embryonic cell cycle (Han et al., 2017). Here we extend that work by showing that the fertilisation induced rise in ROS levels are sustained throughout early embryonic development. To assess the possible roles for these sustained increased ROS levels during mesoderm formation, we treated blastula stage embryos with several antioxidants, including N-acetyl-cysteine (NAC) and MCI-186, and we found that attenuating ROS levels disrupted several signalling pathways, including PI3K/Akt signalling, nodal signalling and canonical Wnt signalling, and as a consequence, mesoderm formation, patterning and morphogenesis were severely impaired. Addition of the oxidants, H_2_O_2_ and menadione, partially restored the impaired signalling caused by the antioxidants. Interestingly, not all signalling pathways were dependent on elevated ROS levels during embryogenesis. This report is, to the best of our knowledge, the first to show that elevated, yet non-pathological levels of ROS play critical roles during vertebrate mesoderm formation.

## Materials and Methods

### Imaging and detection of H_2_O_2_

Transgenic F1 *Xenopus laevis* females expressing HyPer maternally were used (Love et al., 2011). Imaging was performed as previously described (Han et al., 2017).

### Chemical and genetic manipulations

N-acetyl-L-cysteine (NAC), sodium acetate (NaAc), DMSO and H_2_O_2_ were purchased from Sigma. Injections were performed as previously described (Gatherer and Woodland, 1996). A 1 M stock of MCI-186 (Cayman Chemicals) was made in DMSO. A 4 mM stock of Menadione (Sigma) was made in DMSO. Synthetic mRNA was transcribed from pCS2+ derived DNA templates, using mMESSAGE mMACHINE SP6 kit (Ambion).

### Xenopus oocytes manipulation

Stage VI oocytes were manually defolliculated from ovaries and injected with 20 ng HyPer RNA, and incubated at 16 °C for 48 hours in OR2 medium (H. B. Peng, 1991) before use. 2 μM Progesterone was added to the oocytes around 36 hours after RNA injection, to induce oocyte maturation.

### Western blot analysis

Embryos (st.10.5-11) were frozen on dry ice and transferred to minus 80 °C until use. After homogenisation, the equivalent of 1 embryo lysate was loaded onto 8% SDS-PAGE for gel electrophoresis, and transferred to PVDF membrane. The primary antibodies used are: anti-phospho-Akt Ser473 (1:1000, Cell Signalling 4051); anti-Akt (1:1000, Cell signalling 9272); anti-phospho-ERK 1/2 monoclonal antibody (1:10,000, Sigma M7802); anti-Erk (1:1000, Cell signalling 9102); anti-phospho-Smad1/5/8 (1:1000, Cell Signalling 9511); anti-phospho-Smad2 (1:1000, Millipore, 05-953) and anti-α-tubulin antibody: (1:100,000, Sigma 9026). Secondary antibodies were sourced from Dako (UK): anti-rabbit HRP-conjugated (1:40,000, P0448) and anti-mouse HRP-conjugated (1: 100,000, P0447).

### RT-qPCR

Embryos (st.10.5-11) kept in RNAlater were homogenised with a pestle within 24 hours and used for RNA extraction using an RNeasy Mini Kit (QIAGEN). The cDNA synthesis was performed with the High-Capacity RNA-to-cDNA Synthesis Kit (Applied Biosystems) following manufacturer’s instruction. Real-time PCR was performed using SYBR Green PCR Master Mix (Applied Biosystems) and StepOne+ machine. Primers were designed and tested for melting curve profiles prior to experiments. Expression values were generated using the ΔΔCT method and *rpl8* was used as an endogenous control. The following primers were used: rpl8-forward: CACGTGTCCGTGGTGTGGCT; rpl8-reverse: CGACCAGCTGGGGCATCTCT; *bra-forward:* GGGACCCCACCGAGAAGGAG and bra-reverse: GGATCCAGGCCCGACATGCT.

### TOPFlash Assay

Plasmids TOPFlash (100 pg) (Veeman et al., 2003) and pTK-Renilla (50 pg, endogenous control) were co-injected with 50 pg *β-gal* or *wnt8* mRNA into 1-2 cell stage embryos. These embryos were re-injected with 10 nl of 1M NAC or NaAc (final concentration in the embryo around 10 mM), or incubated with 1 mM H_2_O_2_ or 4 μM menadione at st. 8.5 and collected for luciferase analysis at st. 10.5/11. Ten embryos from each group were collected and homogenised with 200 μL 1X PLB, and centrifuged at 4 °C, 16,000 rcf for 10 minutes to separate clear lysate from precipitated yolk. 50 μL clear lysate was aliquot into 96-well plate and analysed with the DLR system for luciferase activity (Promega E1910). M50 TOPFlash and pTK-Renilla is a gift from Randy Moon and Christof Niehrs, respectively. pSP64T-wnt8 originated from Moon lab is a gift from John Gurdon.

### Whole-mount in situ hybridisation

Embryos (st.10.5-11) were fixed and whole-mount in situ hybridisation was performed as previously described (Harland, 1991). Plasmid DNAs were linearized with Bgl II for *in vitro* transcription using T7 RNA polymerase (Roche) with digoxigenin RNA labelling mix (Roche). BM purple (Roche) was used for chromogenic detection.

### Statistical analyses

Statistical analyses were performed using GraphPad Prism^TM^ software. Details are described in each figure legend.

## Results

### An oxidative burst of H_2_O_2_ induced at fertilisation is sustained throughout the early embryogenesis of Xenopus laevis

We have established a method for visualizing H_2_O_2_ in *Xenopus* oocytes and early embryos *in vivo* by employing a transgenic reporter line, which expresses a genetic sensor H_2_O_2_, named Hyper, maternally (Han et al., 2017; Love et al., 2013; 2011). HyPer consists of a prokaryotic OxyR domain and is especially sensitive to hydrogen peroxide (H_2_O_2_) over other ROS. Once oxidised, it induces a reversible conformational change, which can be marked and reflected by two excitations of fluorescence. Thus, we can easily monitor H_2_O_2_ levels *in vivo* by calculating the ratio of HyPer-oxidised and HyPer-reduced (Belousov et al., 2006).

Using this resource, we have previously shown that the H_2_O_2_ levels increased following fertilisation (Han et al., 2017). We next sought to determine whether the fertilisation-induced H_2_O_2_ production was transient or sustained throughout development. Interestingly, we found that the elevated levels of H_2_O_2_ were sustained throughout the early stages of development until the neurula stages, when their levels started to decrease, until reaching a low level at the tadpole stages, less than one half the initial ROS level present in unfertilized eggs (Fig. 1A and B).

**Fig. 1.**
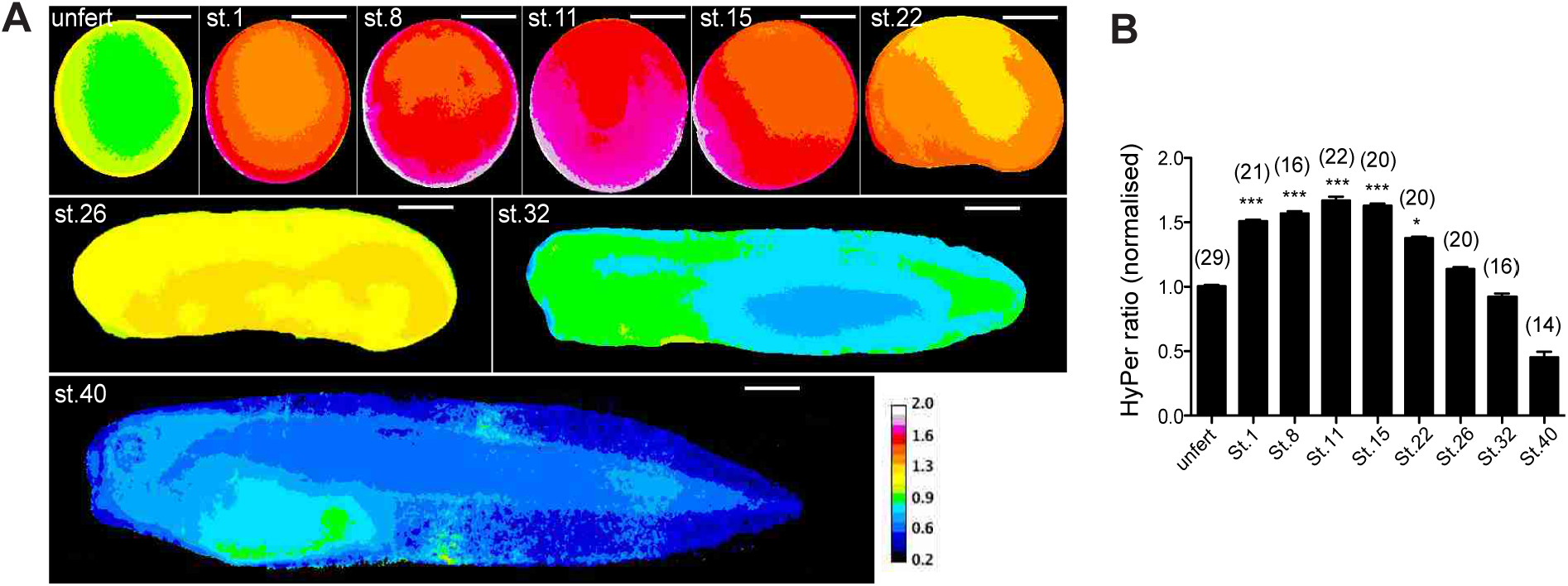
A burst of ROS production triggered by fertilisation is sustained throughout the early embryogenesis. (A) Representative Hyper ratio images of single Xenopus transgenic embryos from an unfertilised egg to the tadpole stage. Panels for St. 26, 32 and 40 were stitched together from multiple images. (B) Quantification of normalised HyPer ratio during embryonic development of X. *laevis.* n number is displayed on the top of each bar. One-Way ANOVA was used for statistical analysis. Error bars represent S.E.M. The asterisks indicate statistically differences at p< 0.05 (*), p< 0.01 (**) and p< 0.001 (***). Scale bar equals 200 micron.

### Lowering embryonic ROS impairs mesoderm formation

To determine the role for the elevated ROS levels during early embryogenesis, we next sought to determine the effect of treating embryos with the antioxidants, N-acetyl-cysteine (NAC) and MCI-186. MCI-186, also referred to as Edavarone, is a novel hydroxyl radical scavenger (Aoyama et al., 2008; Song et al., 2008) and NAC is a precursor of glutathione, and has been shown to reduce oxidative stress *in vivo* (Kerksick and Willoughby, 2005; Shimamoto et al., 2011). Interestingly, it has been shown that injection of NAC into the blastocoel of mid-blastula stage embryos could cause a severe developmental defect in X. *laevis* embryos (Gatherer and Woodland, 1996). However, neither the effect of NAC on ROS nor the molecular mechanisms underlying the developmental defects of NAC on *Xenopus* embryos have been fully investigated. Since we sought to use NAC and MCI-186 as means of decreasing intracellular ROS levels in embryos, we first confirmed that NAC injection and MCI-186 treatment at the blastula stages resulted in attenuated ROS levels at the gastrula and neurula stages (Fig. 2A and B; Supplementary movie). Sodium acetate (NaAc) and DMSO were used as controls for NAC and MCI-186 respectively. Consistent with a previous report (Gatherer and Woodland, 1996), injection of NAC at the blastula stage resulted in severe developmental defects, including enlarged cement glands and failure of axis elongation, a phenotype that was also observed in MCI-186 treated embryos (Fig. 2C). Since the cement gland is derived from the ectoderm (Drysdale and Elinson, 1992) and the germ layer primarily responsible for axis elongation is the mesoderm (Wallingford et al., 2002), we then asked whether mesoderm induction was affected in the antioxidants treated embryos. We first examined the effect of antioxidant treatments on the expression of the general mesodermal marker, brachyury *(bra)* (Smith et al., 1991). Both NAC injected and MCI-treated embryos displayed a significant diminution in *bra* expression at the gastrula stage, as assayed by whole-mount *in situ* hybridisation (Fig. 3A) and qPCR (Fig. 3B), demonstrating that NAC and MCI-186 inhibit mesoderm induction and/or maintenance in X. *laevis* embryos. The decreased expression of *bra* in NAC injected embryos was partially rescued by the addition of H_2_O_2_ to the culture media (Fig. 3C).

**Fig. 2.**
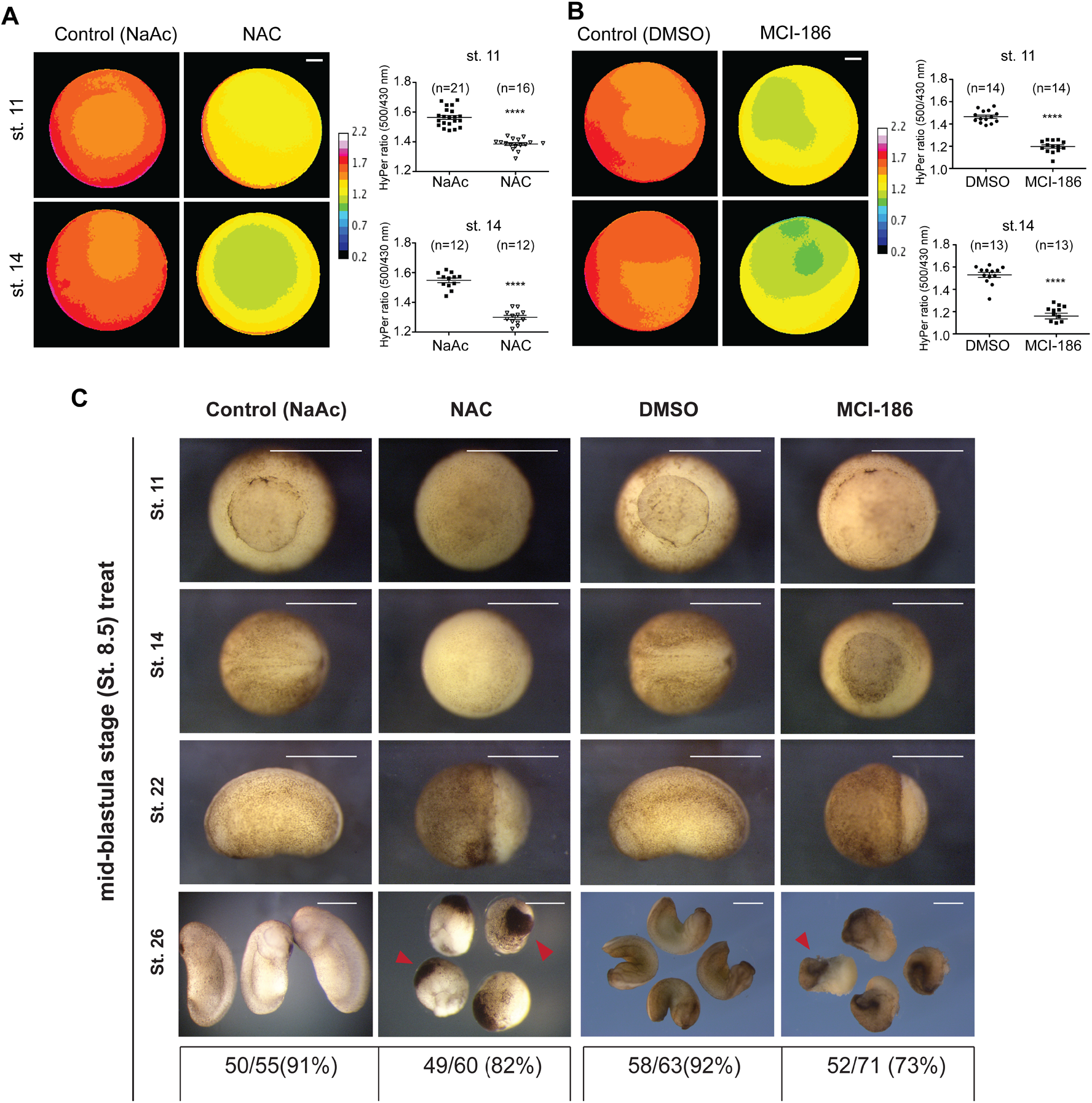
Lowering H_2_O_2_ levels by NAC and MCI-186 induces severe developmental defects in X. *laevis* embryos. (A) HyPer images and quantifications of HyPer ratio in embryos injected with 10 nl of 1 M NAC or 1 M NaAc (final concentration of 10 mM) at the mid-blastula stage (St 8). (B) HyPer images and quantifications HyPer ratio in embryos incubated in 0.1% DMSO or 1 mM MCI-186 from the mid-blastula stage (St 8) and allowed to develop to the indicated stages. Mann-Whitney test, **** denotes P<0.0001. Experiments were repeated three times. (C) Embryonic phenotypes of NAC injected and MCI-186 treated embryos. Numbers of embryos exhibiting phenotypes out of the total number of injected/treated embryos are listed at the bottom. Red arrows indicate enlarged cement glands.

**Fig. 3.**
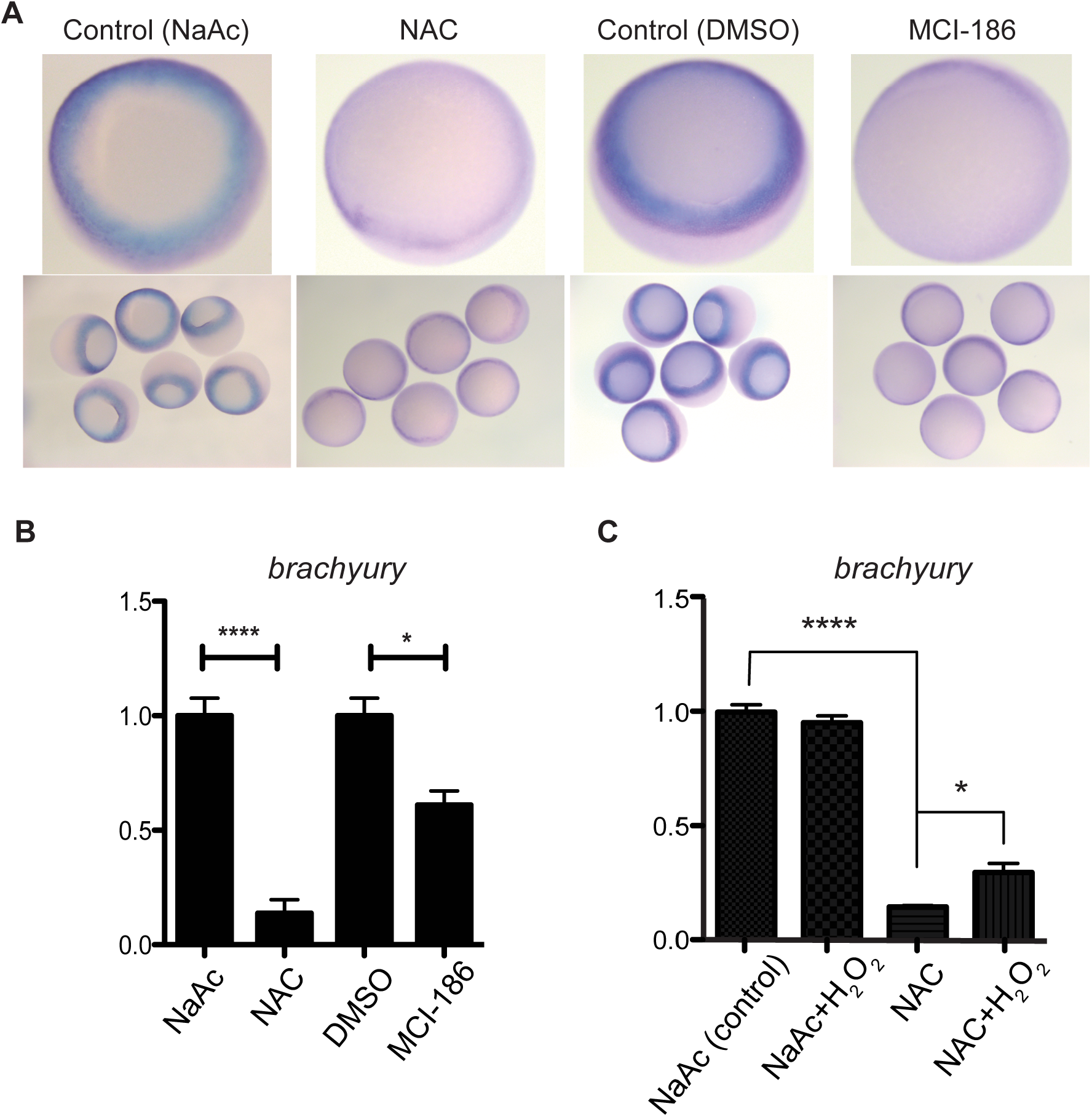
Treatment with the antioxidants, NAC and MCI-186, impairs mesoderm formation. (A) *Xenopus bra* expression in NAC injected / MCI-186 treated embryos was assessed by whole-mount *in situ* hybridisation. Mid-blastula stage (St 8) embryos were injected with 10 nl of 1 M NAC or 1 M NaAc (final concentration of 10 mM) or incubated with in 0.1% DMSO or 1 mM MCI-186 and allowed to develop to the mid-gastrula stage (St 11). (B) *bra* expression from mid-gastrula stage embryos (St 11) was quantified by Real-time PCR (qPCR) following injection of NAC or incubation in MCI-186 as above, *n*=3, analysed by unpaired t-test. (C) Partial rescue of *bra* expression in NAC injected embryos by addition of 1 mM H_2_O_2_ was found in qPCR, n=3, analysed by unpaired t-test. Error bars indicate S.E.M and * denotes P<0.05, ****denotes P<0.0001.

### Elevated ROS levels are required for PI3K/Akt and Nodal signalling

As lowering intracellular ROS levels resulted in the loss of mesoderm induction and/or maintenance, we next sought to identify the underlying molecular mechanisms responsible for the requirement for sustained ROS levels during mesoderm formation. FGF signalling has long been recognised as an essential signalling pathway for mesoderm induction and/or maintenance (Amaya et al., 1991; Dorey and Amaya, 2010; Kroll and Amaya, 1996) and previous work has shown that the downstream effector, PI3K/Akt, participates in mesoderm formation (Carballada et al., 2001). Thus, we first examined whether PI3K/Akt signalling was activated normally in NAC injected or MCI-186 treated embryos. By western blot analysis using an antibody specific for the activated form of Akt, we found that NAC/MCI-186 treatments dramatically reduced phosphorylated Akt (pAkt) levels, but did not alter total Akt levels (Fig. 4A i), suggesting that attenuating ROS levels inhibited Akt activation, without affecting Akt protein levels.

**Fig. 4.**
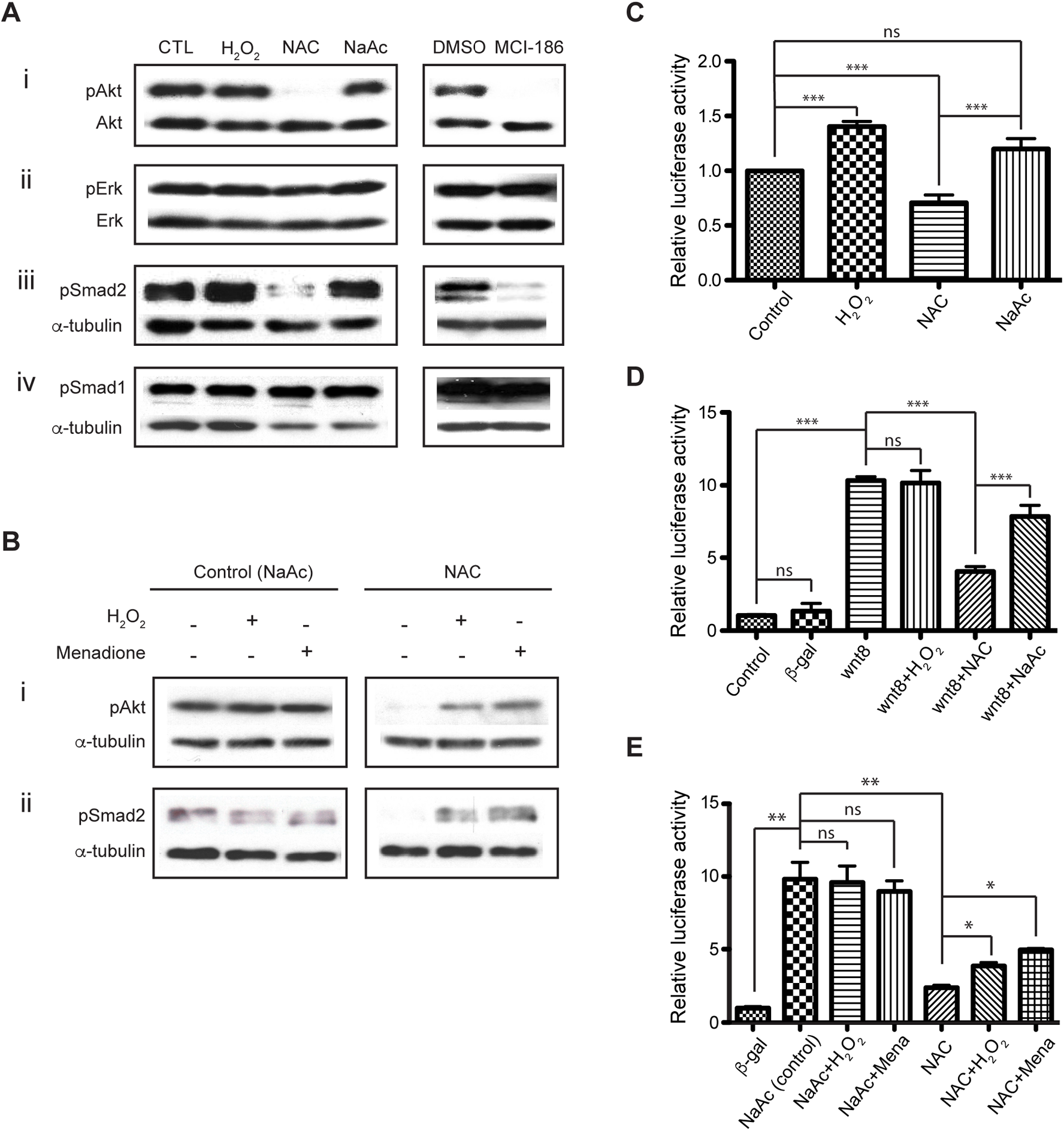
ROS regulate PI3K/Akt, TGF-β and Wnt/β-Catenin signalling. (A) Representative western blots showing that the inhibition of ROS production induces loss of Akt (i) and Smad2 (iii) phosphorylation, but not Erk (ii) and Smad1 (iv) phosphorylation. Mid-blastula stage (St 8) embryos were treated with 1 mM H_2_O_2_, injected with 10 nl of 1 M NAC or 1 M NaAc (final concentration of 10 mM) or incubated with in 0.1% DMSO or 1 mM MCI-186 and collected for analysis at the mid-gastrula stage (St. 11), with antibodies specific to pAkt, pan-Akt, pErk, pan-Erk, pSmad1, pSmad2 and α-tubulin. Experiments were repeated three times. (B) Phosphorylation of Akt and Smad2 were recovered after addition of H_2_O_2_ (1 mM) or menadione (4 μM), to the culture medium. (C) H_2_O_2_ enhanced embryonic Wnt/β-catenin signaling. Embryos injected with TOPFlash and pTK-Renilla plasmids were incubated with 1 mM H_2_O_2_ or injected with NAC or NaAc (control) at the mid-blastula stage (St 8). Ten embryos of each group were collected at the gastrula stage for luciferase analysis. (D) ROS inhibition by NAC attenuates Wnt/β-catenin signalling. Embryos injected with TOPFlash and pTK-Renilla plasmids were co-injected with 50 pg of *wnt8* or *β-gal* (control) mRNA, incubated with 1 mM H_2_O_2_ or injected with NAC or NaAc (control) at st. 8 and collected for luciferase analysis at the mid-gastrula stage (St 11). (E) Partial rescue of Wnt/β-catenin signalling was observed in NAC injected embryos after application of 1 mM H_2_O_2_ or 4 μM menadione. The luciferase assays represent results from at least three independent experiments. Mann-Whitney test was used. Error bars denote S.E.M, The asterisks indicate statistically differences at p< 0.05 (*), p< 0.01 (**) and p< 0.001 (***).

Importantly, we found that addition of exogenous H_2_O_2_ or the oxidant, menadione (Criddle et al., 2006), restored Akt activation in the NAC injected embryos (Fig. 4B i), demonstrating that elevated ROS levels are sufficient to activate PI3K/Akt signalling during *Xenopus* embryogenesis.

Since Erk/MAPK pathway acts in concert with the PI3K/Akt pathway during *Xenopus* mesoderm specification (Carballada et al., 2001; Nie and Chang, 2007), we next asked whether Erk/MAPK signalling was also affected in the antioxidant treated embryos. We found that neither the active, double phosphorylated Erk (dpErk) nor the total Erk protein levels were affected by antioxidant treatments, as evidenced by Western blot analyses (Fig. 4A ii). Thus, our data suggests that, while PI3K signalling is sensitive to ROS levels in gastrula stage embryos, Erk/MAPK signalling is not.

Given that TGF-β family growth factors are also implicated in mesoderm induction and patterning during embryogenesis (Faure et al., 2000; Henry et al., 1996; Kimelman and Kirschner, 1987; Nishimatsu and Thomsen, 1998), we also asked whether antioxidant treatments might affect TGF-β/Nodal/BMP signalling. We found that NAC injected and MCI-186 treated embryos had impaired TGF-β/Nodal signalling, as assessed by assaying for phosphorylated Smad2 (pSmad2) levels (Fig.4A iii). In contrast, however, BMP signalling, as assessed by assaying phosphorylated Smad1 (pSmad1) levels, was not affected by the antioxidant treatments (Fig. 4A iv). It is also worth noting that the loss of pSmad2 levels caused by NAC treatment was also rescued by exogenous H_2_O_2_ or menadione treatments (Fig. 4B ii), suggesting that elevated ROS levels are sufficient to promote TGF-β/Nodal signalling during *Xenopus* embryogenesis as well.

In summary treatment with the antioxidants, NAC and MCI-186, attenuated PI3K/Akt and nodal signalling, but did not affect ERK/MAPK nor BMP signalling.

### Embryonic ROS modulate Wnt/β-catenin signalling

Another important signalling pathway for mesoderm specification is the Wnt/β-catenin signalling pathway, which is required for axis formation and anterior-posterior patterning in *Xenopus* embryos (Hikasa and Sokol, 2013; Huelsken et al., 2000; Larabell et al., 1997). Furthermore, Funato and colleagues showed that addition of H_2_O_2_ can enhance Wnt/β-catenin signalling in mammalian cells *in vitro* (Funato et al., 2006). We then wondered whether early embryonic ROS could similarly regulate Wnt/β-catenin signalling during mesoderm formation. To address this, we carried out a series of experiments on gastrula stage embryos using the TOPFlash reporter assay system, which relies on the quantification of luciferase expression under the transcriptional control of several Wnt/β-catenin regulated TCF/LEF elements (Veeman et al., 2003). We found that embryos treated with H_2_O_2_ from the blastula stage (st. 8.5) showed a significant increase in endogenous Wnt/β-catenin signalling activation at the gastrula stage, relative to control embryos (Fig. 4C). In contrast, gastrula embryos previously injected with NAC displayed significantly lowered endogenous Wnt/β-catenin signalling, relative to control embryos (Fig. 4C). These findings suggest that H_2_O_2_ is capable of hyperactivating endogenous Wnt/β-catenin signalling in embryos, and that endogenous Wnt/β-catenin signalling is dependent on elevated ROS levels in embryos. To determine whether hyper-activated Wnt/β-catenin signalling was also sensitive to ROS levels, we injected *in vitro* transcribed *wnt8* mRNA in embryos and asked whether Wnt/β-catenin signalling activated by Wnt8 overexpression was also sensitive to attenuated ROS levels, following NAC injection. We found that, while H_2_O_2_ was unable to increase Wnt/β-catenin signalling above that seen following Wnt8 overexpression alone, the level of activation achieved by Wnt8 overexpression was significantly attenuated following NAC injection, but not NaAc injection (Fig. 4D). We also found that incubating NAC injected embryos with either exogenous H_2_O_2_ or menadione partially restored Wnt/β-catenin signalling (Fig. 4E). These results suggest that elevated ROS levels facilitate Wnt/β-catenin signalling in *Xenopus* embryos.

## Discussion

The first half of the 20^th^ century marked the primetime of metabolism, as it was during this time when the biochemical pathways underpinning intermediary metabolism were first discovered. It is notable that work on various aspects of intermediary metabolism led to twelve Nobel Prizes in either Physiology and Medicine or Chemistry between 1902 and 1964. Although attention during that time focused primary on the discovery of the enzymatic pathways responsible for anabolic and catabolic pathways, some attention during that time was also devoted to potential roles for metabolism or metabolic signalling in development, as exemplified by Otto Warburg’s pioneering work showing increase in oxygen consumption following fertilisation in sea urchin zygotes (Shearer, 1922; Warburg, 1908). However, the scientist in the first half of the 20^th^ century who most explored the link between metabolism, in particular redox states, and development is Charles Manning Child (Blackstone, 2006). Child’s interests focused largely upon metabolic gradients, most readily measured by redox gradients, in embryos using redox sensitive dyes (Child, 1942). With the advent of genetics and molecular biology in the second half of the 20^th^ century, Child’s pioneering work on metabolic / redox signalling in development quickly became overshadowed by mechanistic studies of development using molecular approaches. However, in the past decade there has been a resurging interest on the potential roles for redox signalling in development.

The work we present here follows upon the pioneering classical work of Warburg, Childs, and others, in that we show that fertilisation results in a burst of ROS production in *Xenopus* embryos, which is sustained throughout early embryogenesis. Furthermore, we show that lowering ROS levels following treatments with the antioxidants, NAC and MCI-186, impairs mesoderm formation, suggesting elevated ROS levels play an essential role during mesoderm formation. Through gain and loss of function analyses, we demonstrated that elevated ROS levels promotes PI3K/Akt, TGF-β/Nodal and Wnt/β-catenin signalling during *Xenopus* early embryogenesis, thus explaining the defects in mesoderm formation after lowering ROS levels in early embryos via treatments with the antioxidants. Although previous studies suggested that ROS could fine-tune several signalling pathways to regulate cellular processes, those studies have largely been done using *in vitro* systems. Using *Xenopus* early embryos as an *in vivo* model system, we provide evidence that elevated ROS levels, likely via redox signalling, play an essential role during a normal physiological system; i.e. vertebrate embryonic development.

The underlying mechanisms by which sustained, elevated ROS levels promotes PI3K, nodal and Wnt/β-catenin signalling, whether directly or indirectly, require further investigation. However, the most likely mechanism by which H_2_O_2_ mediates its effects on growth factor signalling is through its ability to modulate the structure and function of proteins through reversible cysteine oxidation (Forman, 2016; Klomsiri et al., 2011). An example of a redox sensitive protein involved in PI3K signalling is PTEN, which when oxidised by H_2_O_2_ at its catalytic site (Cys^124^), is rendered inactive (Lee et al., 2002). Thus, by inhibiting PTEN activity, H_2_O_2_ promotes PI3K/Akt signalling. In our embryos, which have attenuated ROS levels following antioxidant treatments, PI3K/Akt signalling is lost, possibly due to hyperactive PTEN activity. This finding would also suggest that PI3K/Akt signalling during early embryogenesis is facilitated by increased ROS levels, which attenuates PTEN activity. Since PI3K/Akt has also been shown to have a regulatory role in canonical Wnt signalling via inhibition of glycogen synthase kinase 3β (GSK-3β) in *Xenopus* embryos (Y. Peng et al., 2004), it is also possible that reduced Akt signalling activates GSK-3β, resulting in inhibition of zygotic Wnt/β-Catenin signalling, which leads to dorsalisation of embryo at the gastrula stage.

We also observed a down-regulation of Wnt/β-catenin signalling in gastrula stage embryos after lowering intracellular ROS levels. This finding is consistent with our previous work, showing that inhibiting sustained ROS activation following tail amputation also leads to decreased Wnt/β-catenin signalling (Love et al., 2013). Furthermore, Funato et al. reported that H_2_O_2_ can promote Wnt/β-catenin signalling by catalysing the oxidation of nucleoredoxin (Nrx), a thioredoxin (Trx) related protein, which results in the downregulation of Wnt/β-catenin signalling (Funato et al., 2006). The authors showed that, while oxidised Nrx does not bind to Dishevelled (Dvl), reduced Nrx does. Thus, in the presence of ROS, oxidised Nrx does not inhibit Wnt/β-catenin signalling, but in the absence of ROS, Nrx binds Dvl, resulting in the degradation of β-catenin and inactivation of Wnt/β-catenin signalling (Funato et al., 2006). Thus, ROS may facilitate mesoderm induction/axis specification by modulating PI3K/Akt and Wnt/β-Catenin signalling in either serial and/or parallel pathways.

In our study, we also found that nodal signalling in gastrula embryos is dependent on elevated ROS levels. It was previously shown that mitochondrial ROS/H_2_O_2_ production plays an essential role in localised nodal expression in sea urchins, and thus elevated ROS/H_2_O_2_ levels plays a critical role in the establishment of the oral-aboral axis in sea urchin embryos (Coffman et al., 2009). Given that the expression of several nodal genes are dependent on Wnt/β-catenin signalling in *Xenopus* embryos (Houston, 2017), it is perhaps not surprising that nodal signalling is impaired in embryos with reduced ROS levels, given the requirement for elevated ROS levels in embryos for proper activation of Wnt/β-catenin signalling (this study). However, it is also notable that previous work has suggested that elevated ROS levels can be involved in the release of active TGF-β (Liu and Desai, 2015), thus it is perhaps important to assess in the future whether elevated ROS levels may also have a role in activating Nodal ligands, and not just their expression during embryogenesis.

We previously found that tail amputation results in sustained elevated ROS levels, which are necessary for tail regeneration in tadpoles (Love et al., 2013). We also found in that study that inhibiting ROS production during tail regeneration resulted in impaired Wnt/β-catenin signalling and *fgf20* expression, which are both required for tail regeneration. A role for elevated ROS levels during appendage and tissue regeneration has been described in a variety of organisms, including zebrafish, planarians, fruit flies, geckos and rats (Bai et al., 2015; Gauron et al., 2013; Khan et al., 2017; Pirotte et al., 2015; Santabárbara-Ruiz et al., 2015; Zhang et al., 2016). The finding that early embryos are also associated with sustained elevated ROS levels and that these elevated ROS levels also regulate growth factor signalling in development, suggests that, successful appendage regeneration is necessarily accompanied by a return to an embryonic-like redox state, which is permissive for growth factor signalling. It is, thus, interesting to speculate whether sustained ROS levels, which create a permissive environment for both embryonic development and appendage regeneration could be harnessed in order to promote tissue regeneration in species with poor regenerative capacities, such as humans.

## Acknowledgements

We thank Christoph Niehrs for the pTK-Renilla plasmid. We extend gratitude to the University of Manchester Bioimaging Facility for guidance with imaging. This work was supported by a Wellcome Trust Program Grant (E.A.), an MRC project grant (EA), two PhD studentships from The Healing Foundation (Y.H., Y.C) and a grant from The Healing Foundation (N.R.L., E.A.).

## Supplementary Movie. Reduced ROS levels were observed after injection of NAC

Live imaging of HyPerYFP during embryogenesis with injection of NAC and NaAc (control).

## References

Amaya, E., Musci, T.J., Kirschner, M.W., 1991. Expression of a dominant negative mutant of the FGF receptor disrupts mesoderm formation in Xenopus embryos. Cell 66, 257–270.

Aoyama, T., Hida, K., Kuroda, S., Seki, T., Yano, S., Shichinohe, H., Iwasaki, Y., 2008. Edaravone (MCI-186) scavenges reactive oxygen species and ameliorates tissue damage in the murine spinal cord injury model. Neurol. Med. Chir. (Tokyo) 48, 539–45– discussion 545.

Bai, H., Zhang, W., Qin, X.-J., Zhang, T., Wu, H., Liu, J.-Z., Hai, C.-X., 2015. Hydrogen Peroxide Modulates the Proliferation/Quiescence Switch in the Liver During Embryonic Development and Posthepatectomy Regeneration. Antioxid. Redox Signal. 22, 921–937. doi:10.1089/ars.2014.5960

Bedard, K., Krause, K.-H., 2007. The NOX family of ROS-generating NADPH oxidases: physiology and pathophysiology. Physiol. Rev. 87, 245–313. doi: 10.1152/physrev.00044.2005

Belousov, V.V., Fradkov, A.F., Lukyanov, K.A., Staroverov, D.B., Shakhbazov, K.S., Terskikh, A.V., Lukyanov, S., 2006. Genetically encoded fluorescent indicator for intracellular hydrogen peroxide. Nat. Methods 3, 281–286. doi:10.1038/nmeth866

Blackstone, N.W., 2006. Charles Manning Child (1869-1954): the past, present, and future of metabolic signaling. J. Exp. Zool. 306B, 1–7. doi:10.1002/jez.b.21085

Brand, M.D., 2016. Mitochondrial generation of superoxide and hydrogen peroxide as the source of mitochondrial redox signaling. Free Radic. Biol. Med. 100, 14–31. doi: 10.1016/j.freeradbiomed.2016.04.001

Carballada, R., Yasuo, H., Lemaire, P., 2001. Phosphatidylinositol-3 kinase acts in parallel to the ERK MAP kinase in the FGF pathway during Xenopus mesoderm induction. Development 128, 35–44.

Child, C.M., 1942. Oxidation-reduction patterns in amphibian and teleost development. Proc. Natl. Acad. Sci. U.S.A 28, 339–343.

Cho, S.-H., Lee, C.-H., Ahn, Y., Kim, H., Kim, H., Ahn, C.-Y., Yang, K.-S., Lee, S.-R., 2004. Redox regulation of PTEN and protein tyrosine phosphatases in H(2)O(2) mediated cell signaling. FEBS Lett. 560, 7–13.

Coffman, J.A., Coluccio, A., Planchart, A., Robertson, A.J., 2009. Oral-aboral axis specification in the sea urchin embryo III. Role of mitochondrial redox signaling via H2O2. Dev Biol 330, 123–130. doi:10.1016/j.ydbio.2009.03.017

Coffman, J.A., Wessels, A., DeSchiffart, C., Rydlizky, K., 2014. Oral-aboral axis specification in the sea urchin embryo, IV: hypoxia radializes embryos by preventing the initial spatialization of nodal activity. Dev Biol 386, 302–307. doi:10.1016/j.ydbio.2013.12.035

Criddle, D.N., Gillies, S., Baumgartner-Wilson, H.K., Jaffar, M., Chinje, E.C., Passmore, S., Chvanov, M., Barrow, S., Gerasimenko, O.V., Tepikin, A.V., Sutton, R., Petersen, O.H., 2006. Menadione-induced reactive oxygen species generation via redox cycling promotes apoptosis of murine pancreatic acinar cells. J. Biol. Chem. 281, 40485–40492. doi:10.1074/jbc.M607704200

Dorey, K., Amaya, E., 2010. FGF signalling: diverse roles during early vertebrate embryogenesis. Development 137, 3731–3742. doi:10.1242/dev.037689

Drysdale, T.A., Elinson, R.P., 1992. Cell Migration and Induction in the Development of the Surface Ectodermal Pattern of the Xenopus laevis Tadpole. Development, Growth & Differentiation 34, 51–59. doi:10.1111/j.1440-169X.1992.00051.x

Faure, S., Lee, M.A., Keller, T., Dijke, ten, P., Whitman, M., 2000. Endogenous patterns of TGFbeta superfamily signaling during early Xenopus development. Development 127, 2917–2931.

Foerder, C.A., Klebanoff, S.J., Shapiro, B.M., 1978. Hydrogen peroxide production, chemiluminescence, and the respiratory burst of fertilization: interrelated events in early sea urchin development. Proc. Natl. Acad. Sci. U.S.A. 75, 3183–3187.

Forman, H.J., 2016. Redox signaling: An evolution from free radicals to aging. Free Radic. Biol. Med. 97, 398–407. doi:10.1016/j.freeradbiomed.2016.07.003

Funato, Y., Michiue, T., Asashima, M., Miki, H., 2006. The thioredoxin-related redox-regulating protein nucleoredoxin inhibits Wnt-beta-catenin signalling through dishevelled. Nat. Cell Biol. 8, 501–508. doi:10.1038/ncb1405

Gatherer, D., Woodland, H.R., 1996. N-acetyl-cysteine causes a late re-specification of the anteroposterior axis in the Xenopus embryo. Dev. Dyn. 205, 395–409. doi: 10.1002/(SICI)1097-0177(199604)205:4<395::AID-AJA4>3.0.CO;2-D

Gauron, C., Meda, F., Dupont, E., Albadri, S., Quenech’Du, N., Ipendey, E., Volovitch, M., Del Bene, F., Joliot, A., Rampon, C., Vriz, S., 2016. Hydrogen peroxide (H2O2) controls axon pathfinding during zebrafish development. Dev Biol 414, 133–141. doi:10.1016/j.ydbio.2016.05.004

Gauron, C., Rampon, C., Bouzaffour, M., Ipendey, E., Teillon, J., Volovitch, M., Vriz, S., 2013. Sustained production of ROS triggers compensatory proliferation and is required for regeneration to proceed. Sci Rep 3, 2084. doi:10.1038/srep02084

Han, Y., Ishibashi, S., Iglesias-Gonzalez, J., Chen, Y., Love, N.R., Amaya, E., 2017. Ca^2+^-induced mitochondrial ROS regulate the early embryonic cell cycle. bioRxiv 223123.

Harland, R.M., 1991. In situ hybridization: an improved whole-mount method for Xenopus embryos. Methods Cell Biol. 36, 685–695.

Heinecke, J.W., Shapiro, B.M., 1989. Respiratory burst oxidase of fertilization. Proc. Natl. Acad. Sci. U.S.A. 86, 1259–1263.

Henry, G.L., Brivanlou, I.H., Kessler, D.S., Hemmati-Brivanlou, A., Melton, D.A., 1996. TGF-beta signals and a pattern in Xenopus laevis endodermal development. Development 122, 1007–1015.

Hikasa, H., Sokol, S.Y., 2013. Wnt signaling in vertebrate axis specification. Cold Spring Harb Perspect Biol 5, a007955–a007955. doi:10.1101/cshperspect.a007955

Houston, D.W., 2017. Vertebrate Axial Patterning: From Egg to Asymmetry. Adv. Exp. Med. Biol. 953, 209–306. doi:10.1007/978-3-319-46095-6_6

Huelsken, J., Vogel, R., Brinkmann, V., Erdmann, B., Birchmeier, C., Birchmeier, W., 2000. Requirement for beta-catenin in anterior-posterior axis formation in mice. J. Cell Biol. 148, 567–578.

Kamata, H., Honda, S.-I., Maeda, S., Chang, L., Hirata, H., Karin, M., 2005. Reactive oxygen species promote TNFalpha-induced death and sustained JNK activation by inhibiting MAP kinase phosphatases. Cell 120, 649–661. doi:10.1016/j.cell.2004.12.041

Kerksick, C., Willoughby, D., 2005. The antioxidant role of glutathione and N-acetyl-cysteine supplements and exercise-induced oxidative stress. J Int Soc Sports Nutr 2, 38–44. doi:10.1186/1550-2783-2-2-38

Khan, S.J., Abidi, S.N.F., Skinner, A., Tian, Y., Smith-Bolton, R.K., 2017. The Drosophila Duox maturation factor is a key component of a positive feedback loop that sustains regeneration signaling. PLoS Genet 13, e1006937. doi: 10.1371/journal.pgen.1006937

Kimelman, D., Kirschner, M., 1987. Synergistic induction of mesoderm by FGF and TGF-beta and the identification of an mRNA coding for FGF in the early Xenopus embryo. Cell 51, 869–877.

Klomsiri, C., Karplus, P.A., Poole, L.B., 2011. Cysteine-based redox switches in enzymes. Antioxid. Redox Signal. 14, 1065–1077. doi:10.1089/ars.2010.3376

Kroll, K.L., Amaya, E., 1996. Transgenic Xenopus embryos from sperm nuclear transplantations reveal FGF signaling requirements during gastrulation. Development 122, 3173–3183.

Larabell, C.A., Torres, M., Rowning, B.A., Yost, C., Miller, J.R., Wu, M., Kimelman, D., Moon, R.T., 1997. Establishment of the dorso-ventral axis in Xenopus embryos is presaged by early asymmetries in beta-catenin that are modulated by the Wnt signaling pathway. J. Cell Biol. 136, 1123–1136.

Lee, S.-R., Yang, K.-S., Kwon, J., Lee, C., Jeong, W., Rhee, S.G., 2002. Reversible inactivation of the tumor suppressor PTEN by H2O2. Journal of Biological Chemistry 277, 20336–20342. doi:10.1074/jbc.M111899200

Liu, R.-M., Desai, L.P., 2015. Reciprocal regulation of TGF-β and reactive oxygen species: A perverse cycle for fibrosis. Redox Biology 6, 565–577. doi: 10.1016/j.redox.2015.09.009

Love, N.R., Chen, Y., Ishibashi, S., Kritsiligkou, P., Lea, R., Koh, Y., Gallop, J.L., Dorey, K., Amaya, E., 2013. Amputation-induced reactive oxygen species are required for successful Xenopus tadpole tail regeneration. Nat. Cell Biol. 15, 222–228. doi: 10.1038/ncb2659

Love, N.R., Thuret, R., Chen, Y., Ishibashi, S., Sabherwal, N., Paredes, R., Alves-Silva, J., Dorey, K., Noble, A.M., Guille, M.J., Sasai, Y., Papalopulu, N., Amaya, E., 2011. pTransgenesis: a cross-species, modular transgenesis resource. Development 138, 5451–5458. doi:10.1242/dev.066498

Nie, S., Chang, C., 2007. PI3K and Erk MAPK mediate ErbB signaling in Xenopus gastrulation. Mech. Dev. 124, 657–667. doi:10.1016/j.mod.2007.07.005

Nishimatsu, S., Thomsen, G.H., 1998. Ventral mesoderm induction and patterning by bone morphogenetic protein heterodimers in Xenopus embryos. Mech. Dev. 74, 75–88.

Peng, H.B., 1991. Xenopus laevis: Practical uses in cell and molecular biology. Solutions and protocols. Methods Cell Biol. 36, 657–662.

Peng, Y., Jiang, B.-H., Yang, P.-H., Cao, Z., Shi, X., Lin, M.C.M., He, M.-L., Kung, H.-F., 2004. Phosphatidylinositol 3-kinase signaling is involved in neurogenesis during Xenopus embryonic development. Journal of Biological Chemistry 279, 28509–28514. doi: 10.1074/jbc.M402294200

Pirotte, N., Stevens, A.-S., Fraguas, S., Plusquin, M., Van Roten, A., Van Belleghem, F., Paesen, R., Ameloot, M., Cebrià, F., Artois, T., Smeets, K., 2015. Reactive Oxygen Species in Planarian Regeneration: An Upstream Necessity for Correct Patterning and Brain Formation. Oxid Med Cell Longev 2015, 392476–19. doi:10.1155/2015/392476

Ray, P.D., Huang, B.-W., Tsuji, Y., 2012. Reactive oxygen species (ROS) homeostasis and redox regulation in cellular signaling. Cell. Signal. 24, 981–990. doi: 10.1016/j.cellsig.2012.01.008

Robinson, K.A., Stewart, C.A., Pye, Q.N., Nguyen, X., Kenney, L., Salzman, S., Floyd, R.A., Hensley, K., 1999. Redox-sensitive protein phosphatase activity regulates the phosphorylation state of p38 protein kinase in primary astrocyte culture. J. Neurosci. Res. 55, 724–732. doi: 10.1002/(SICI)1097-4547(19990315)55:6<724::AID-JNR7>3.0.CO;2-9

Santabárbara-Ruiz, P., López-Santillán, M., Martínez-Rodríguez, I., Binagui-Casas, A., Pérez, L., Milán, M., Corominas, M., Serras, F., 2015. ROS-Induced JNK and p38 Signaling Is Required for Unpaired Cytokine Activation during Drosophila Regeneration. PLoS Genet 11, e1005595–26. doi:10.1371/journal.pgen.1005595

Shearer, C., 1922. On the Oxidation Processes of the Echinoderm Egg During Fertilisation. Proceedings of the Royal Society B: Biological Sciences 93, 213–229. doi: 10.1098/rspb.1922.0016

Shimamoto, K., Hayashi, H., Taniai, E., Morita, R., Imaoka, M., Ishii, Y., Suzuki, K., Shibutani, M., Mitsumori, K., 2011. Antioxidant N-acetyl-L-cysteine (NAC) supplementation reduces reactive oxygen species (ROS)-mediated hepatocellular tumor promotion of indole-3-carbinol (I3C) in rats. J Toxicol Sci 36, 775–786.

Smith, J.C., Price, B.M., Green, J.B., Weigel, D., Herrmann, B.G., 1991. Expression of a Xenopus homolog of Brachyury (T) is an immediate-early response to mesoderm induction. Cell 67, 79–87.

Song, Y., Gong, Y.-Y., Xie, Z.-G., Li, C.-H., Gu, Q., Wu, X.-W., 2008. Edaravone (MCI-186), a free radical scavenger, attenuates retinal ischemia/reperfusion injury in rats. Acta Pharmacol. Sin. 29, 823–828. doi:10.1111/j.1745-7254.2008.00822.x

Tseng, W.-C., Munisha, M., Gutierrez, J.B., Dougan, S.T., 2017. Establishment of the Vertebrate Germ Layers. Adv. Exp. Med. Biol. 953, 307–381. doi:10.1007/978-3-319-46095-6_7

Veeman, M.T., Slusarski, D.C., Kaykas, A., Louie, S.H., Moon, R.T., 2003. Zebrafish prickle, a modulator of noncanonical Wnt/Fz signaling, regulates gastrulation movements. Current Biology 13, 680–685.

Wallingford, J.B., Fraser, S.E., Harland, R.M., 2002. Convergent extension: the molecular control of polarized cell movement during embryonic development. Dev. Cell 2, 695–706.

Warburg, O., 1908. Beobachtungen über die Oxydationsprozesse im Seeigelei. Hoppe-Seyler’s Zeitschrift für physiologische Chemie 57, 1–16. doi: 10.1515/bchm2.1908.57.1-2.1

Wong, J.L., Créton, R., Wessel, G.M., 2004. The oxidative burst at fertilization is dependent upon activation of the dual oxidase Udx1. Dev. Cell 7, 801–814. doi:10.1016/j.devcel.2004.10.014

Zhang, Q., Wang, Y., Man, L., Zhu, Z., Bai, X., Wei, S., Liu, Y., Liu, M., Wang, X., Gu, X., Wang, Y., 2016. Reactive oxygen species generated from skeletal muscles are required for gecko tail regeneration. Sci Rep 6, 20752. doi:10.1038/srep20752

